# Time-resolved cryoEM using Spotiton

**DOI:** 10.1101/2020.04.20.050518

**Authors:** Venkata P. Dandey, William C. Budell, Hui Wei, Daija Bobe, Kashyap Maruthi, Mykhailo Kopylov, Edward Eng, Peter A. Kahn, Jenny E. Hinshaw, Nidhi Kundu, Crina M. Nimigean, Chen Fan, Nattakan Sukomon, Seth Darst, Ruth Saecker, James Chen, Brandon Malone, Clinton S. Potter, Bridget Carragher

**Affiliations:** The National Resource for Automated Molecular Microscopy, Simons Electron Microscopy Center, New York Structural Biology Center, 89 Convent Ave, New York, NY, 10027, USA; Engineering Arts LLC, Arizona, USA; Laboratory of Cell and Molecular Biology, National Institute of Diabetes and Digestive and Kidney Diseases, NIH, Bethesda, MD, 20892, USA; Department of Anesthesiology, Weill Cornell Medical College, New York, NY, 10065 USA; Laboratory of Molecular Biophysics, The Rockefeller University, New York, NY 10065, USA; Department of Biochemistry and Molecular Biophysics, Columbia University, New York, NY 10032, USA

**Keywords:** time-resolved, short-lived molecular states, cryoEM vitrification, piezo dispensing, nanowire grids

## Abstract

We present an approach for preparing cryoEM grids to study short-lived molecular states. Using piezo electric dispensing, two independent streams of ~50 pL sample drops are deposited within 10 ms of each other onto a nanowire EM grid surface, and the mixing reaction stops when the grid is vitrified in liquid ethane, on the order of ~100 ms later. We demonstrate the utility of this approach for four biological systems where short-lived states are of high interest.

---

Cryo electron microscopy (cryoEM) has the distinct advantage of being able to capture a wide variety of conformational states of macromolecules in solution. Changes in conformational states can be triggered by a variety of biological reactions. For example, by adding a ligand to an enzyme, mixing together components of a multimolecular machine, or by adding energy in the form of ATP or GTP. These conformational changes are often transient but can be trapped by vitrification of the sample at specific time points following the initiation of the reaction and then imaged using electron microscopy, a process which has been loosely referred to as “time-resolved cryoEM”^1^ when applied to the study of a conformational process occurring on the millisecond time scale. For much slower processes, standard vitrification or negative staining at fixed time points (on the order of seconds to minutes) serves the same goal^2, 3^.

Possibly the earliest approach to time-resolved cryoEM was by Berriman and Unwin^4^ where one reactant in the form of small droplets was sprayed onto another in the form of a thin aqueous film supported by an EM grid substrate. Fast timing was achieved by locating the sprayer just above the cryogen cup into which the grid was plunged - thereby stopping the reaction - and by spraying onto a grid that was already moving at high speed towards the cryogen. The potential of this approach to reveal conformational changes induced by ligands on fast-acting molecules, such as ion channels, was demonstrated by high-resolution images obtained of acetylcholine receptor tubes onto which acetylcholine had been sprayed^5^. A little later, the group of Howard White described an updated computer-controlled device and demonstrated its efficacy in observing the interaction of myosin sprayed onto actin tubes^6^. The method is generally described as “spraying-mixing” and can achieve time resolutions as low as 2 ms^7^. A disadvantage of this approach is that mixing is not uniform across the grid and in order to identify specific areas where the sprayed droplets mix with the standing solution, some kind of fiducial marker must be present in the sprayed solution. In another approach to time-resolved cryoEM, conformational changes in the proton pump bacteriorhodopsin were observed by exposing crystals of bacteriorhodopsin, sitting on an EM grid, to variable periods of light illumination followed by rapid freezing in liquid ethane^8^. This method also provides millisecond control over the timing between the light-induced action and the trapped conformational state but is only suitable for photo-active samples.

While these methods engendered a lot of interest, difficulties with practical implementation resulted in very few further publications using time-resolved cryoEM until more recently, when an alternative “mixing-spraying” approach was developed by the group of Joachim Frank^9,10^. Their device was based on the design described by Howard White^11^, but mixed the two samples prior to spraying small droplets of the mixture onto a dry grid plunging rapidly towards a cryogen. Mixing and spraying were achieved using a microfabricated device that enabled very fast mixing, using chaotic advection^12^, followed by a fixed reaction chamber length to control the reaction time prior to pneumatic spraying. The system has been used in a variety of time-resolved experiments to study the mechanics of ribosomes^13, 14, 15, 16^ and is capable of reaction times of as low as ~10 ms. A recent paper^17^ described an update that allows for either mixing-spraying or spraying-mixing. Applying this approach to proteins other than the ribosome may present difficulties, including the need for a fairly large (~30 μL) volume of protein sample for each grid. In addition, the thickness of the vitreous ice layer varies with droplet size and the spreading of the droplet on the grid, potentially reducing the efficiency of data collection.

We have developed a new approach to the spraying-mixing method based on the Spotiton robot^18, 19^ that uses a piezo dispensing tip to apply a stream of ~50 pL droplets onto a nanowire (“self-wicking”) grid^20^ as it rapidly speeds past on its way to vitrification in a liquid cryogen. This method produces a stripe of ice of fairly uniform thickness across each grid, which is often sufficient to acquire enough data for a high-resolution map. The fast spot-to-plunge time also has some value in ameliorating the deleterious effects of the air-water interface and has been used to prepare grids for a wide variety of protein samples^21–26^. By adding a second piezo dispensing tip to the device, we can deliver a second stream of droplets onto the first stream within 10 ms of it being deposited. The two sample volumes mix on the grid as the bulk volume is wicked away and spread out to a thin film by the capillary action of the nanowires. We describe the method and demonstrate its efficacy and value for four biological systems where short-lived states are of high interest: (i) binding of ribosomal subunits; (ii) binding of promoter DNA to RNA polymerase; (iii) binding of Ca^2+^ to a potassium channel followed by a conformational change; (iv) conformational rearrangements of dynamin lipid tubes driven by GTP hydrolysis.

Volumes on the order of 50 pL have been shown to mix completely within ~10 ms when brought together in mid-air just before colliding with a surface^27^ and thus good mixing of the drops on the nanowire surface is expected. Nevertheless, as a first proof of principle to validate the basic operation of time-resolved Spotiton, we mixed two abundantly available, well behaved, and well understood test samples, apoferritin and 70S ribosomes. As shown in Figure 1, we dispensed the two samples (Table S5) onto the grid by setting the second stream of sample droplets to be initially overlapping the first stream at the leading edge of the grid and then separated from it by 1-3 squares towards the end of sample deposition nearer the grid’s trailing edge. In this way, we are able to provide the unmixed control experiments at the same time as the mixed. As observed in the images in Figure 1, in the non-overlapping regions we see apoferritin and 70S ribosomes in high concentrations and well distributed in the vitreous ice of the individual separated stripes, whereas in the overlapping area, we observe both particles well mixed.

**Figure 1:**
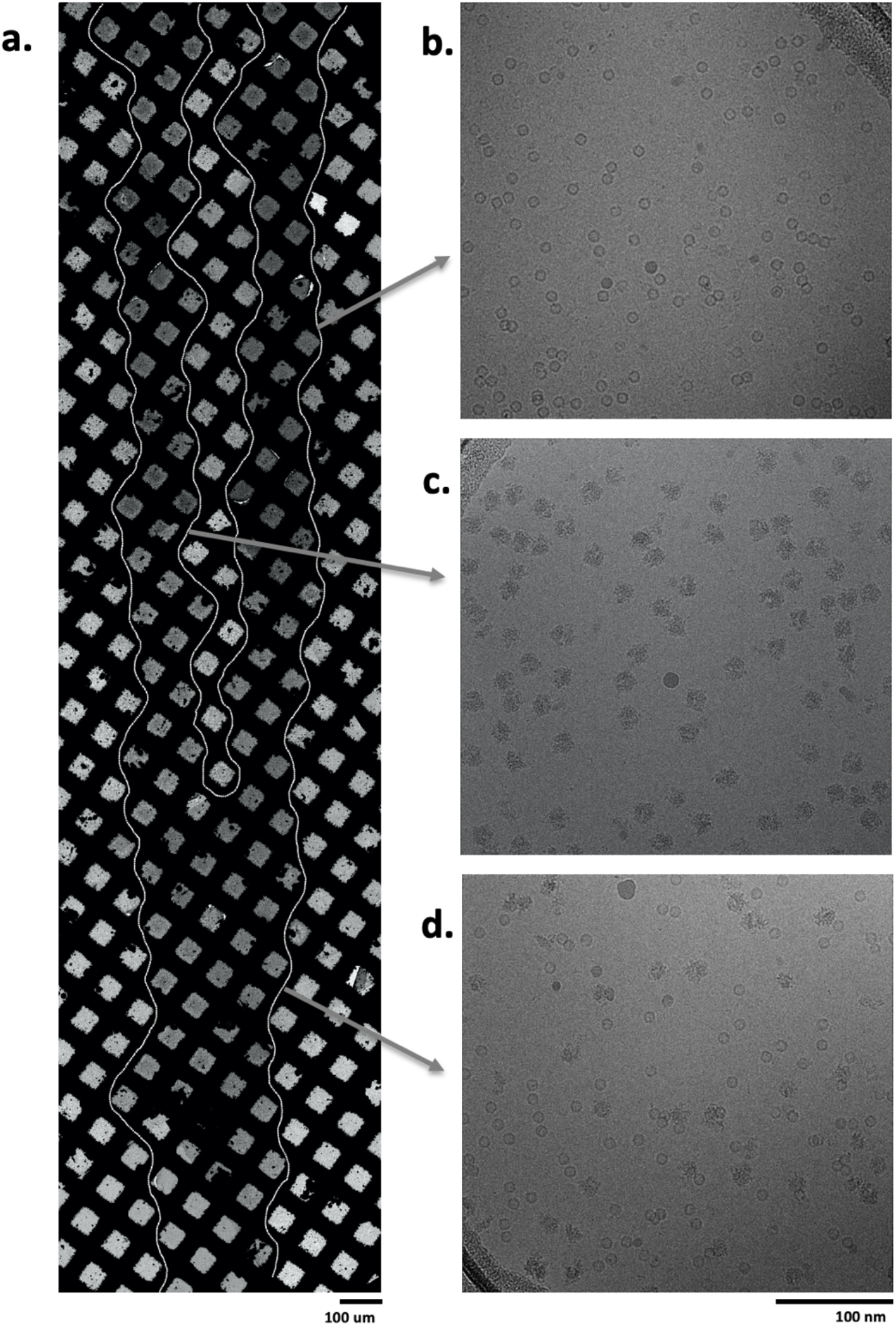
Apoferritin and 70S ribosomes were used as a proof of concept to illustrate mixing on the nanowire grids. (a) Overview of the vitrified grid showing the sample streams merged at the leading edge {bottom) and separated at the trailing edge (top). Squares containing vitrified ice are indicatedby a white outline. Example micrographs obtained from the indicated regions show either only {b) apoferritin or (c) 70S ribosomes or (d) a mix of both samples.

Early work on mixing-spraying demonstrated that separated 30S and 50S ribosomal subunits could associate into 70S ribosomes^10^. In Figure 2a we show an example of an image of the overlapping area of grid where these two sample have been mixed. Control experiments of the individual samples showed populations of 30S or 50S subunits (plus a small percentage of 50S dimer particles), and no evidence of 70S complexes (Figure S2c, d). The mixed sample contained 30S, 50S, 50S dimers plus about 20% assembled 70S ribosomes (Figure 2a); this particle count was estimated by picking all particles in the field of view and then sorting in 2D and 3D to arrive at a reconstructed map of the 70S ribosome at a resolution of 4.8 Å (Figure S2a,b). We note that previous work^14^ observed ~40% assembled 70S ribosomes using a mixing-spraying device followed by a 140 ms delay in a reaction chamber. This difference in assembly states is not surprising as the rate of interactions between subunits is expected to be much slower via diffusion within droplets that mix on the grid than by the mixing that occurs by chaotic advection in the device described in the earlier study. We also note that the previous study used a higher concentration of 30S subunits, twice that of 50S subunits, whereas the data shown here used a 1:1 ratio of 30S to 50S subunits.

**Figure 2:**
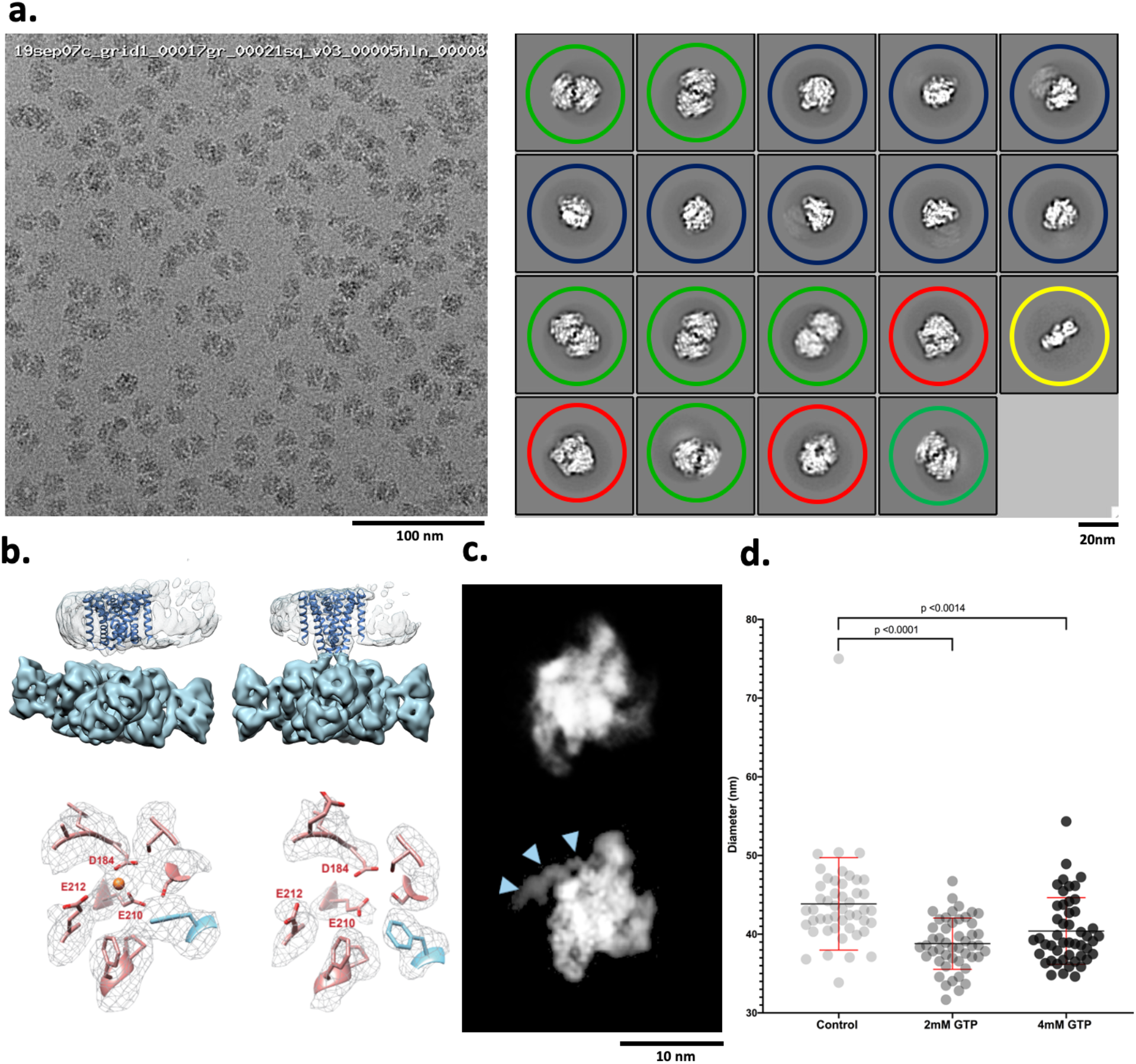
Four examples of biological systems where time-resolved cryoEM provides answers. (a) *left:* Representative micrograph from the mixed region of a Spotiton prepared grid shows 30S and 50S ribosomal subunits and 70S complexes; *right:* corresponding 2D classes show particles representing a population of 30S (yellow), 50S (blue), 50S-50S dimers (green), and 70S (red). ~20% of particles were reconstructed to a 70S complex at a resolution of 4.75 A (see Methods and Figure S2). (b) 3D maps generated from MthK in the presence *(top left)* or absence *(top right*) of calcium showing clear differences in the overall conformation of the channel. The bottom row shows one of the three Ca^2+^ binding sites in MthK either occupied in the case of a mixing experiment *(left)* or vacant as with the MthK only control *(right*). (c) Representative class averages of RNAP alone *(top)* or mixed with promoter DNA *(bottom)* showing DNA clearly bound (blue arrowheads). (d) Measured diameters (mean ± SD) of dynamin-decorated tubes without (control: 43.85, ± 5.86, n=48) and with GTP (2 mM: 38.80, ± 3.2, n=48; 4 mM: 40.40, ± 4.23, n=48,). Student’s t test p values are shown.

An obviously compelling use of time-resolved cryoEM is to observe changes in an ion channel during the early stages of its interaction with a ligand. We examined the conformational changes in a calcium-gated potassium channel (MthK^28–30^) upon interacting with calcium ions. When reconstituted into liposomes, MthK was observed to activate within milliseconds after application of saturating Ca^2+^ concentrations and then to inactivate slowly with a time constant of ~2 s^39^. This slow inactivation is caused by the 17 N-terminal amino acids entering the pore from the intracellular side and obstructing ion permeation in a ball-and-chain-like manner^47^. As predicted from functional assays, single-particle cryoEM analyses of MthK channels in the absence and presence of Ca^2+^ yield structures of closed and inactivated channels, respectively, with the inactivated structure prominently displaying the N-terminus lodged inside the pore^47^. The obvious prediction from these results is that freezing MthK channels on cryoEM grids within ~100 ms of Ca^2+^ application, after the channel opens but before it inactivates, will reveal the structure of an open channel and other intermediates, completing the picture of the gating cycle. Using time-resolved Spotiton to mix MthK and Ca^2+^, we observed that at ~150 ms, the major class resulting from 3D classification has the TM domain in the nanodisc tilted with respect to the large extra-membranous density (Figure 2b, top). This pronounced tilt is one of the hallmarks of a Ca^2+^-bound open MthK state and is different from the closed MthK structure obtained in the absence of Ca^2+^, which displays little to no tilt^47^ (Figure 2b, top). Analysis of the large ligand binding domain (RCK gating ring) alone from all classes revealed that the major conformation of the gating ring indeed corresponds to that of the open MthK structure^31, 47^ (Figure S3b). In addition, densities for Ca^2+^ ions were observed at all known binding sites in the open MthK structure (Figure 2b, bottom and Figure S3a). While the data is not yet sufficient to identify if the channel is open or inactivated - as the density for the transmembrane domains within the nanodiscs is weak - these results indicate that Ca^2+^ not only successfully mixed and bound to MthK, but also managed to induce a conformational change to an activated state.

RNA synthesis by all DNA-dependent RNA polymerases (RNAP) is a tightly regulated dynamic process involving large-scale conformational changes in both the enzyme and DNA. Mechanistic investigations of the formation of the transcriptionally-competent open complex, Rpo, by bacterial RNAP have defined a multi-step pathway where a series of intermediates appear and disappear on the subsecond to seconds time scale^32, 33^. While “kinetically-significant” intermediates in RPo formation were identified decades ago^34^, their transient nature has prevented atomic resolution structural characterization. As a consequence, the conformational changes involved in their isomerizations and how they are targeted by regulatory factors remain largely unknown. While the use of temperature or other variables have historically been used to trap intermediates at equilibrium in solution, the ultimate goal is to capture structural snapshots of them as they interconvert in time. To this end, we examined DNA opening by *E. coli* RNAP holoenzyme at the λP_R_ promoter using time-resolved Spotiton. The kinetics of RPo formation at λP_R_ have been extensively characterized, allowing predictions of intermediate populations as a function of time and solution conditions^32^. Within the ~150 ms time of mixing and freezing only the earliest intermediates are predicted to be present. The 2D class averages from this experiment show DNA bound to RNAP in a conformation consistent with promoter recognition (Figure 2c). Future experiments that vary time, solution conditions, and promoter sequence combined with 3D classification strategies are anticipated to reveal the on-pathway nucleation and propagation of the transcription bubble.

Finally, we looked at the dynamics of dynamin at the ~150 ms timescale. During GTP hydrolysis, dynamin constricts and pinches off invaginating clathrin-coated vesicles. Previous results have shown dynamin constricts the membrane within seconds and during this process the helical parameters transform from a 1-start to a 2-start helix^35, 36^. However, the rate and mechanism of how the dynamin polymer constricts and rearranges during GTP hydrolysis remain unknown. Using time-resolved Spotiton to mix pre-formed dynamin tubes with GTP, we observed that at ~150 ms, a high percentage of the dynamin decorated tubes were constricted (i.e. the lumen of the lipid bilayer was reduced) to 39 nm upon mixing with 2 mM GTP compared to 44 nm for untreated controls (Figure 2d). Upon mixing with 4 mM GTP, the dynamin polymer becomes disordered, constricts, and disassembles from the lipid bilayer (Figure 2d and S4). This work provides the first clues to the initial steps that lead to dynamin-mediated membrane constriction and fission. We expect further analysis incorporating a decreasing range of GTP concentrations will trap the reaction at the slowest step, allowing changes in the dynamin organization during early fission events to be observed.

These four biological cases represent a range of short-lived molecular states of high interest and demonstrate that samples can be successfully mixed on a grid and rapidly vitrified within ~100 ms to trap intermediates present at this time scale. This method uses very small quantities of material and is applicable to mixing together any two, or potentially more, samples to allow the capture of short-lived molecular states that appear between 50-500 ms after an initial interaction.

## Methods

### Spotiton Instrumentr

The Spotiton system^18, 19^ was upgraded with a second set of identical dispense head components. A second piezo driven electric tip was mounted next to the first head 4.5 mm from the first tip (Figure S1b). The 4.5 mm pitch allows the two tips to simultaneously aspirate sample from two adjacent holder tubes also mounted at a 4.5 mm pitch. The second tip includes a manual fine adjustment screw to allow precise alignment between the two tips in the direction perpendicular to the plunge axis motion.

The second piezo electric tip is driven by an independent electronic drive (DE03 controller) which was added to the system. The plunge axis outputs a series of electrical pulses while plunging (distance between pulses is a parameter set to 0.25 mm). The plunge axis pulse output is tied to the trigger input of both DE03 controllers. Each DE03 controller can be setup to start firing its respective tip after a unique number of pulses (configurable by the user) relating to the position of the plunge axis. The fluidics of the second piezo electric tip are attached to a second syringe pump which was added to the system. The syringe pumps allow precise independent sample aspiration and cleaning of the tips.

#### Time resolution

The motion path (136 mm) of a grid prepared by the Spotiton time-resolved system is characterized by three phases: acceleration, constant velocity, and deceleration (Figure S1a). The durations of the acceleration and constant velocity phases are variable and dependent on the rate of acceleration set by the user. The deceleration phase remains fixed and ends when the grid comes to a stop in the liquid ethane cup. The two dispensers spray a defined number of sample droplets whose first contact with the falling grid is separated by a period of time between 3-7 ms, depending on the acceleration rate selected. The first sample thus has a brief opportunity to be wicked away by the capillary action of the nanowires prior to contact by the second sample. (Figure S1b) The mixing time of the two samples prior to vitrification ranges from 130-160 ms but can be reduced to ~90 ms by moving the dispensers to a “low-fire” position ~4 cm closer to the ethane bath (Condition 4 in Table S1). A redesigned instrument would in principle be capable of even shorter mixing times; for example, the commercial version of the Spotiton system, Chameleon (SPT Labtech), is capable of spot-to-plunge times of ~50 ms.

#### Time-resolved Spotiton operation

A typical protocol for operating the time-resolved Spotiton system proceeds as follows. On startup, the system is initialized, and the two three-axis robots used to position the grid-holding tweezers and the dispensing heads are homed. Next, the fluid lines carrying distilled water from an external reservoir to the dispensing tips are flushed several times to remove air bubbles and any residual methanol used to clean the tips after the previous session. Both tips are then fired in view of an inspection camera to confirm successful dispensing and the formation of discrete droplets^19^. Next, a standard (not nanowire) test grid is loaded into the tweezers, lowered into position between the upper camera and the upper dispenser tip, and both are observed for alignment in the live viewer in the main software window. Aided by an integrated image-recognition algorithm, the operator positions the tip along the vertical midline and at the leading (lower) edge where the first dispensed droplets will contact the grid. The second piezo device is locked into position directly beneath the first, but its tip is steerable to allow the second sample to be dispensed either completely overlapping the first sample strip or in a discrete, non-contacting parallel strip. To verify operation, each tip is fired separately on the test grid and video captures from the upper camera are examined to confirm deposition of a liquid stripe. To verify tip alignment (i.e. both stripes are deposited onto the same grid area), both tips are fired simultaneously, and the video capture is examined for the presence of a single overlapping liquid stripe. Once successful two-tip dispensing on the test grid is confirmed, the humidity of the chamber is increased to 80-85% by activating the in-chamber nebulizer. Next, 5 μL of each sample is loaded into the sample holder cups, placed in the humidified chamber, and simultaneously aspirated into the two tips. Successful firing is again confirmed as described above.

The system is now ready to prepare vitrified grids. First, the upper tip is positioned in front of the upper camera as before and a plasma treated, nanowire grid is loaded into the tweezers. To avoid saturating the nanowires with water, thereby reducing the wicking capacity, exposure of the grid to the high humidity environment within the chamber is limited to 10-60 s prior to plunging, depending on the observed performance of the particular batch of nanowire grids being used. During a typical plunge, a grid acceleration of 10 m/s^2^ and a tip firing frequency of 14,750 Hz results in the deposition of ~70 droplets (~4 nl volume) of each sample onto the grid. This is observed as a single thick, opaque band of liquid down the grid in the upper camera video capture that wicks to a thin film that is nearly invisible in the lower camera video capture acquired 49 ms later, just before the grid plunges into the liquid ethane (Figure S1a). To generate control grids with two non-overlapping sample strips separated by several squares, the lower, steerable tip can be adjusted to bring the tips out of alignment, as mentioned above, or more simply, we can change the acceleration of the grid. During our characterization of the system, we noted that when the tips are aligned to form completely overlapping stripes at a set acceleration (e.g. 10 m/s^2^), the stripes can, at other accelerations, become misaligned, e.g. merging of only the leading (8 m/s^2^; see Figure 1a) or trailing (5 m/s^2^) ends, or even deposited as two parallel and completely separated stripes (6 m/s^2^). While we do not fully understand the mechanics of this phenomenon, it is reliably reproducible and can be used to make a control grid with two well separated streams of sample by a simple adjustment of the acceleration.

Compared to the original Spotiton protocols^19^, grid preparation and timing was adjusted to account for wicking of double the usual sample volume. This required optimizing our self-wicking grids^20^ to have longer length and higher density nanowires to create a faster and higher volume wicking area.

### Sample preparation

A series of experiments was performed to first test and verify mixing of protein samples on the grids and then to demonstrate the value of this approach for a variety of biological systems of interest. In general, vitrified grids of mixed samples were prepared as follows. Nanowire grids were freshly plasma cleaned and transferred into the humidity chamber (set to 80-85% humidity) no more than 30 seconds prior to vitrification. 5 μL of each sample was loaded into the two sample holder cups with concentrations as tabulated in Table S5. For control experiments, the second sample was replaced by the carrying buffer as noted. The calculated spot-to-plunge time for all of the grids is 151 ms. Below we briefly describe further details of sample and grid preparation for each of these experiments.

#### Apoferritin and 70S ribosomes

Apoferritin was purchased from Sigma Aldrich, 400 kDa, A3660, 2.3 mg/ml. Protein solution stored in 50% glycerol was exchanged into a cryo compatible buffer (50 mM Tris-Cl [pH 7.6]; 150 mM NaCl) using Amicon Ultra-15 centrifugal filter units (100 kDa cutoff membrane). 70S ribosomes were purchased from New England BioLabs Inc, 2MDa. Protein solution was stored in 20 mM HEPES-KOH [pH 7.6], 10 mM Mg(CH_3_COO)_2_, 30 mM KCl, and 7 mM β-mercaptoethanol after diluting the sample to 1 mg/ml from 33.3 mg/ml.

#### 30S an 50S ribosomal subunits

70S ribosomes are prepared as described in^37^. For subunit purification, 70S ribosomes were exchanged into dissociation buffer (20 mM MES-KOH [pH 6], 600 mM KCl, 8 mM Mg(CH_3_COO)_2_, 1 mg/ml heparin, 0.1 mM PMSF, 0.1 mM benzamidine, and 2 mM DTT) before loading onto a sucrose gradient in the same buffer and centrifuged for 19 hr at 28,500 RPM in the Ti25 rotor. The 50S and 30S subunits were exchanged separately into reassociation buffer (10 mM MES-KOH [pH 6], 10 mM NH_4_CH_3_COO, 40 mM CH_3_COOK, 8 mM Mg(CH_3_COO)_2_, and 2 mM DTT), concentrated to 6 μM, and stored at −80°C after being flash frozen in liquid nitrogen.

#### RNA Polymerase plus promoter DNA

Core RNAP (subunit composition α_2_ββ’ω) was expressed and purified as described^38^. The specificity subunit σ^70^ was expressed and purified as described^38^ with the following modifications: i. a plasmid encoding His(6)- SUMO- σ^70^ was used instead of His(10)-SUMO-σ^70^; ii. cells were grown at 30°C in the presence of 50 μg/mL kanamycin until OD 0.4, then temperature was lowered to 16°C; iii. when the cells reached OD 0.7, 0.1 mM IPTG was added and growth continued for an additional 15 hrs. After harvest by centrifugation and resuspension in lysis buffer^38^, cells were flash frozen in liquid nitrogen and stored overnight at −80°C. Cells were thawed halfway at 22°C, thawed completely on ice, and then lysed in a French press. After lysis, the series of columns and buffers used to purify σ^70^ were as described ^38^. For promoter DNA, a duplex λP_R_ promoter fragment (−85 to +20) was used (Trilink Biotechnologies, San Diego, CA). Top (non-template) strand: ‘5 C GGA ATC GAG GGA TCC TCT AGA GTT GGA TAA ATA TCT AAC ACC GTG CGT GTT GAC TAT TTT ACC TCT GGC GGT GAT AAT GGT TGC ATG TAC TAA GGA GGT TGTA G 3’. Bottom (template-strand): 5’ C TACA ACC TCC TTA GTA CAT GCA ACC ATT ATC ACC GCC AGA GGT AAA ATA GTC AAC ACG CAC GGT GTT AGA TAT TTA TCC AAC TCT AGA GGA TCC CTC GAT TCC G 3’. RNAP holoenzyme was assembled by mixing core with a 3.3 molar excess of σ^70^, incubating for 20 min at 37°C, and buffer exchanging into gel filtration (GF) buffer (40 mM Tris-HCl [pH 8.0], 120 mM KCl, 10 mM MgCl_2_ and 10 mM DTT) using centrifugal filtration (Amicon-Ultra-0.5 m 30K cutoff) at 4°C. Excess σ^70^ was separated from core RNAP on a Superose 6 increase 10/300 GL column (GE Healthcare) equilibrated in GF buffer. The eluted fractions of RNAP were concentrated to 16 mg/ml (centrifugal filtration), aliquoted, flash frozen in liquid nitrogen, and stored at −80°C. The non-template and template strands of λP_R_ promoter DNA were dissolved in annealing buffer (10 mM Tris-HCl [pH 8], 50 mM KCl, 0.1 mM EDTA), mixed in equimolar amounts and incubated in a 95°C heat block for 10 min. The samples were then slow cooled in the heat block to room temperature. Annealed DNA was stored at −80°C. RNAP and DNA aliquots were thawed on ice, diluted to the concentrations reported in Table S5 with GF buffer. N-octyl-β-D-glucopyranoside (Anatrace) was added to 0.35% final just before spraying.

#### Ca^2+^ activated channel MthK

MthK was purified and reconstituted into nanodiscs composed of 3:1 POPE:POPG, following the procedure described in detail previously^39, 47^.

#### Dynamin tubes plus GTP

Liposome formation and dynamin purification: 1,2-dioleoyl-sn-glycero-3-phospho-L-serine (100 l of 5 mg/ml, DOPS, Avanti) was dried and resuspended in 250 μl HCB150 (50 mM HEPES, 150 mM KCl, 2 mM EGTA, 1 mM MgCl_2_, 1 mM TCEP, [pH 7.5]). Unilamellar liposomes were obtained by extruding the mixture 21 times through a 0.4 μm pore-size polycarbonate membrane (Avanti). Recombinant ΔPRD-dynamin 1 was purified from Sf9 insect cells. Briefly, recombinant baculovirus containing the sequence of ΔPRD-dynamin 1 with an N-terminal His-tag, was generated by following Bac-to-Bac Baculovirus Expression System (ThermoFisher Scientific). The suspension cultures of Sf9 were maintained in Sf-900 III serum-free media (SFM, ThermoFisher Scientific) and inoculated with recombinant baculovirus at a cell density of 1.6 × 10^6^ with 1/100 volume of virus/final volume of medium. The cells were grown for 72 h at 27°C, and pelleted by centrifugation at 1000 × g, 5 min, 4°C. The pellet was resuspended in 3 μl modified HSB150 (50 mM HEPES, 150 mM KCl, 5 mM beta-mercaptoethanol,10 mM imidazole, [pH 8.0]) and containing protease inhibitor cocktail (Millipore Sigma). The cells were then lysed by sonication (total time of 8 min with 5 s pulse-on and 15 s pulse-off) followed by high speed centrifugation (20,000 × *g*, 15 min). The supernatant was collected, passed through Ni-NTA beads and the protein was eluted with 150 mM imidazole in modified HSB150. The protein solution was dialyzed in HSB150 overnight and the purity was checked using SDS-PAGE/Coomassie staining. Dynamin decorated tubes were generated by incubating 3 μl of DOPS liposomes with 40 μl of protein (0.8 mg/ml, in 10 mM Tris, 10 mM KCl, 1 mM MgCl_2_, [pH 7.4]) for 2 h.

### Imaging and analysis

Typically, data was acquired using Leginon MS^40^ and micrographs were collected either on a Titan Krios (Thermo Fisher Scientific) with a K2 or K3 BioQuantum counting camera (Gatan, Inc.) operating in counting mode or on a Tecnai F20 equipped with a TVIPS CMOS camera. The nominal magnification, pixel size, exposure time, frame rate, total dose, and defocus range were as shown in Table S6 for each experiment. For all datasets, frames were aligned using MotionCorr2^41^ and CTF was estimated using Ctffind4^42^.

#### 70S association complex, 50S and 30S ribosomal subunits

Particle picking was performed with Gautomatch (http://www.mrc-lmb.cam.ac.uk/kzhang/) and extracted in Relion by 5X binning followed by one round of 2D classification using Relion^48^ to remove false particles. After this first round of 2D classification, classes clearly representing 50S dimers and 30S and 50S particles (Figure 2a) were excluded from further steps of image analysis. After one round of 3D classification, only recognizable 70S particles were selected and reextracted to a pixel size of 2.2 Å for final refinement. A total of 26,402 particles were used for homogeneous 3D refinement in Relion resulting in a 4.8 Å map of the 70S ribosome (Figure S2a and b). For the control experiments, a procedure similar to that described above was used to obtain a total of 12,505 and 3,762 individual 50S and 30S particles, respectively, and 2D classified in Relion (Figures S2c and d).

#### MthK with and without Ca^2+^

Typically, a small set of particles was manually picked, and 2D class averages were calculated using the CL2D algorithm^43^ inside the Appion image processing pipeline^44^. A subset of these classes was used as templates to pick particles for the entire set of micrographs using FindEM^45^. For MthK without Ca^2+^, from 956,882 particles and after several rounds of 2D and *ab initio* classification in Cryosparc^49^. 428,917 particles were used for a final 3D classification and the best class was selected and used for Cryosparc2 non-uniform refinement to generate a structure with an overall resolution of 4.2 Å as shown in top right of Figure 2b.

For MthK with Ca^2+^, a procedure was used similar to that described above. Briefly, 2,158,345 particles were auto picked and used for 2 rounds of 2D classification in Relion3. From these, 849,864 good particles were selected and used for 3D classification in Relion3. The open state class with a highly tilted RCK domain was selected and used for Cryosparc2 non-uniform refinement to generate a structure with an overall resolution of Å as shown in top left of Figure 2b.

For the focused refinement of the RCK domain, signal subtracted particles of both samples were generated with a mask to only include the RCK domain. These particles were used for refinement in Relion3 applying C2 symmetry and the overall resolution is Å for Mthk with Ca^2+^ and 3.5 Å for MthK without Ca^2+^ (Figure S3b).

#### RNAP with and without DNA

Particle picking was performed with Gautomatch and extracted in Relion followed by one round of 2D classification to remove false picks using 2D classification tool in Cryosparc^49^. 167,212 particles of RNAP with λP_R_ promoter DNA and 52,747 particles of RNAP alone are used for another round of 2D classification and the 2D classes with high resolution features were selected (Figure 2c).

#### Dynamin with and without GTP

46, 28, and 100 images were collected for dynamin-decorated tubes mixed with 4 mM GTP, 2 mM GTP and no GTP, respectively. For each condition, the diameters of 48 tubes were measured using Fiji^46^.

---

**Figure S1:**
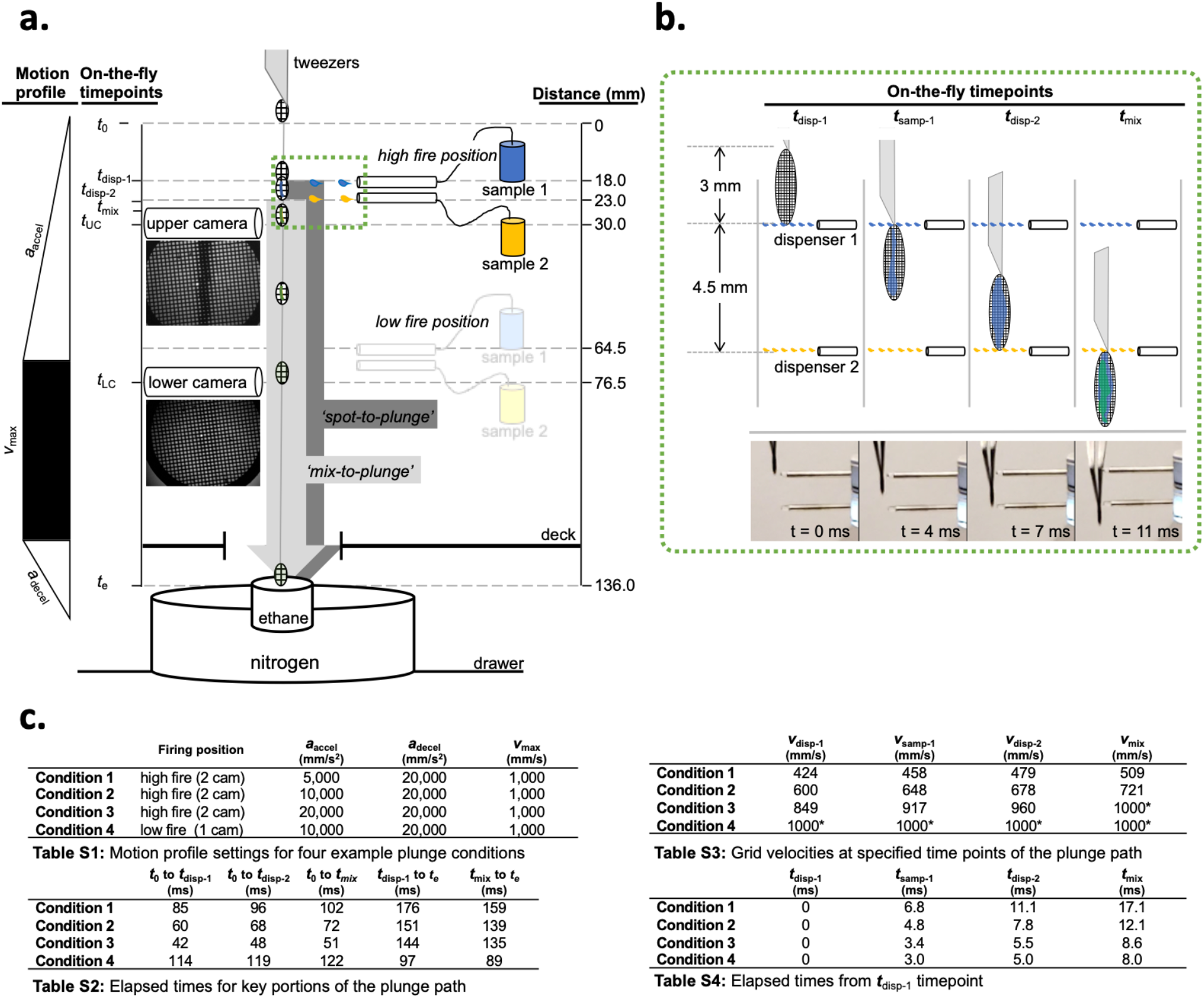
Specifications of time-resolved Spotiton operation. (a) Diagrammatic overview of the distances (fixed) and elapsed times (variable) relevant to spraying and mixing two samples on a moving grid. Simultaneous dispensing of both samples is triggered after the grid plunge begins. Representative images from the upper and lower cameras are shown directly below the illustrations of each. Sample 1 and sample 2 are indicated in blue and yellow, respectively. (b) Magnified view of (green-dashed) boxed area in (a) showing grid and dispensing at specific time-points with corresponding high-speed video captures of the tips and grid below. Elapsed times shown on each image reflect estimates from a video of a grid plunged under Condition 2. Objects in (a) and (b) are not drawn to scale. Tables S1-S4 in (c) show values for the following parameters of a grid plunge as depicted in (a) and (b): *a*_accel_ acceleration rate; *a*_decel_, deceleration rate; *V*_max_, maximum velocity; *t*_0_, plunge start point; *t*_disp-1_, grid leading edge reaches first dispenser; *t*_samp-1_, sample 1 fully applied to grid; *t*_disp-2_, grid leading edge reaches second dispenser; *t*_mix_, samples 1 and 2 fully applied to grid; *t*_UC_, grid reaches upper camera, *t*_Lc_, grid reaches lower camera; *t*_*e*_, grid plunges into ethane. ‘Spot-to­ plunge’ and ‘mix-to-plunge’in (a) reflect the elapsed times from *t*_disp-1_ or *t*_mix_ to *t*_*e*_, respectively.

**Figure S2:**
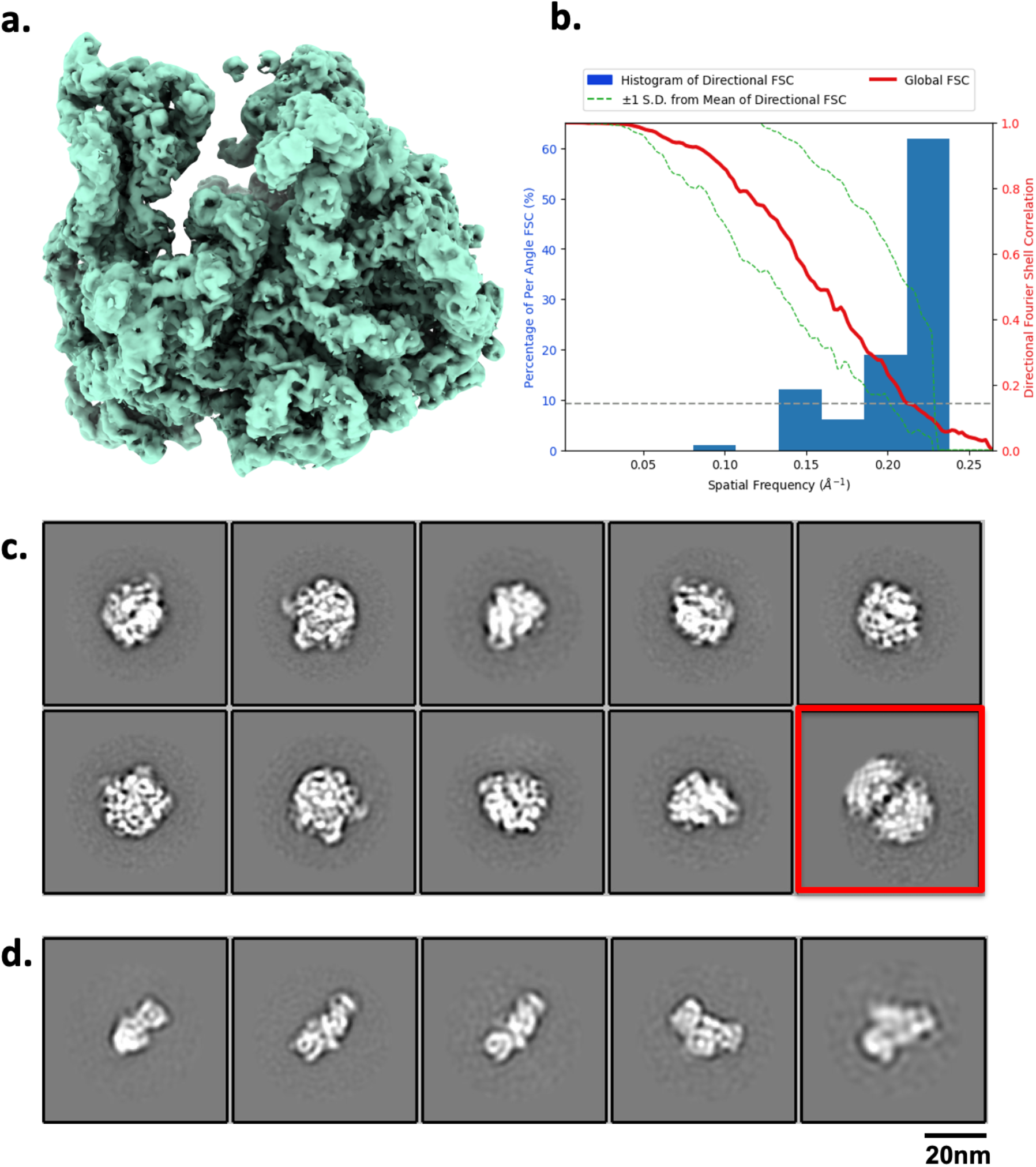
Mixing 30S and 50S ribosomal subunits to form 70S complexes. (a) ~20% of particles present were reconstructed to 70S complex at a resolution of 4.75 A as indicated by FSC_0.5_ (b). (c) 2D classes of 50S ribosomal subunit obtained from the control experiment; 2D class of 50S-50S dimer is shown in red. (d) 2D classes of the 30S ribosomal subunit obtained from the control experiment. Both control experiments show no evidence of 70S ribosomes as observed in the mixed experiment.

**Figure S3:**
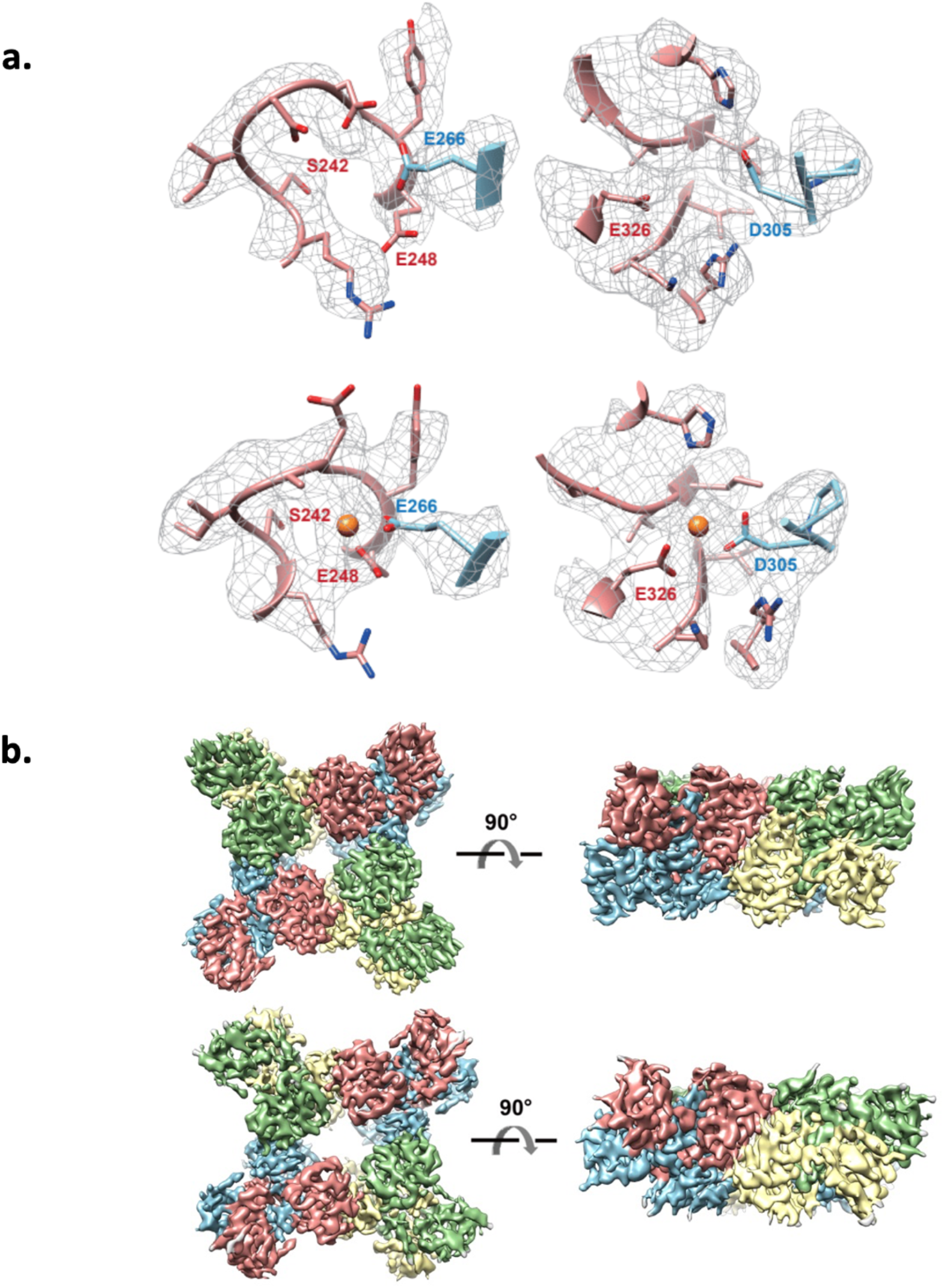
Cryo-EM maps of MthK RCK domain with and without Ca^2+^ (a) The two additional Ca^2+^ binding sites of MthK either vacant from a control experiment (top row) or occupied after mixing with calcium (bottom row). (b) 3D models of MthK RCK domains without (top row) and without (bottom row) Ca^2+^ bound.

**Figure S4:**
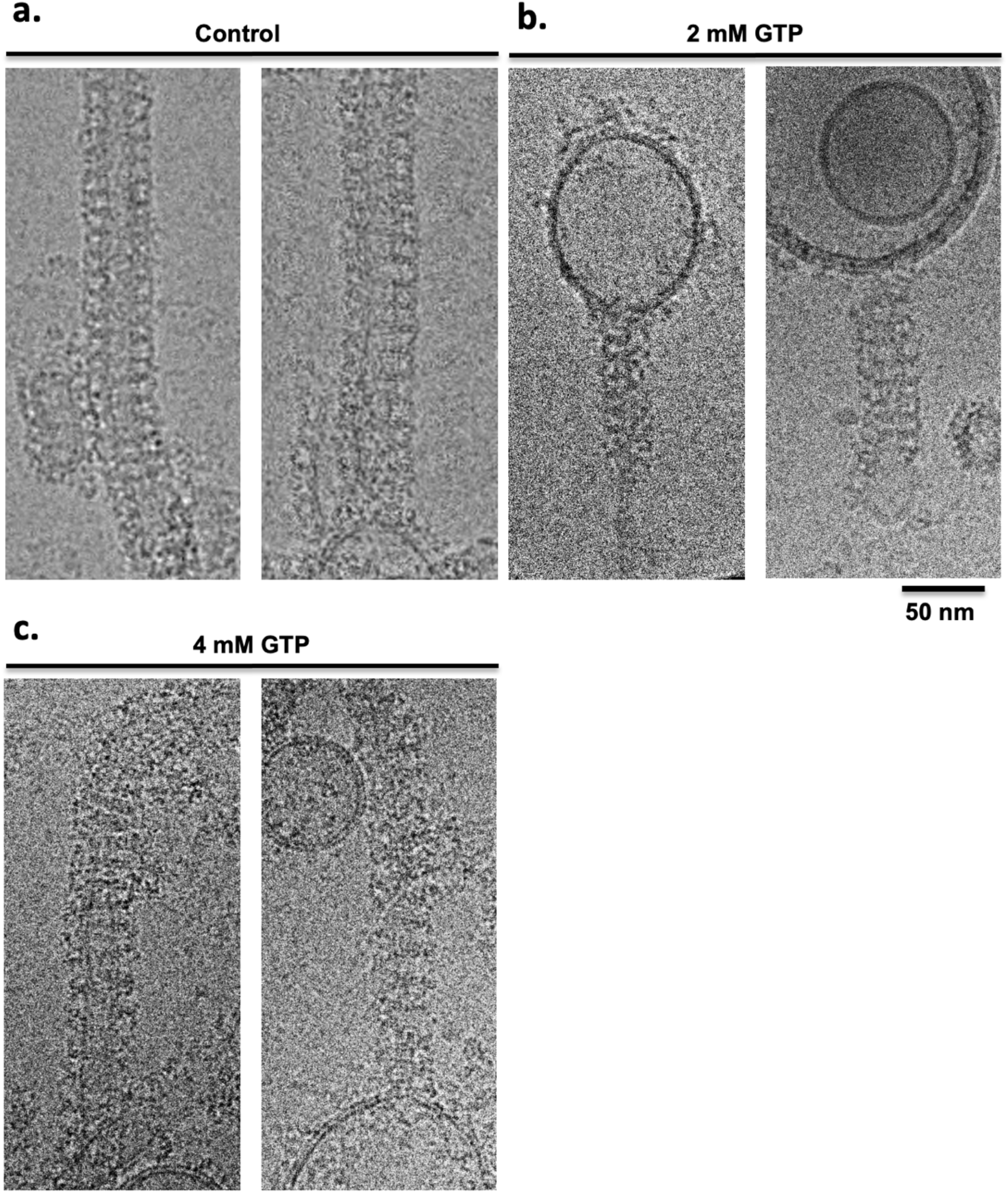
Mixing of GTP with dynamin-decorated lipid tubes results in constriction. Representative cryo-electron micrographs of control dynamin-decorated tubes without GTP (a), with 2 mM GTP (b) and 4mM GTP (c).

**Table S5:** Spotiton sample preparation conditions.

| Experiment | Sample (concentration in dispenser) |
| --- | --- |
| ApoF + 70S mixed | Tip 1 Apoferritin (2.3 mg/ml) |
|  | Tip 2 70S ribosomes (1mg/ml) |
| 50S + 30S mixed | Tip 1 50S (1.4 μM) |
| | Tip 2 30S (1.4 μM) |
| 50S or 30S only | Tip 1 50S or 30S (1.4 μM) |
|  | Tip 2 Buffer |
| MthK + Ca<sup>2+</sup> mixed | Tip 1 MthK (12 mg/ml), Fos8-F (3 mM) |
|  | Tip 2 Ca<sup>2+</sup> (30 mM), Fos8-F (3 mM) |
| MthK only | Tip 1 MthK (12 mg/ml), Fos8-F (3 mM) |
|  | Tip 2 Buffer, Fos8-F (3 mM) |
| RNAP + DNA mixed | Tip 1 DNA (24 μM), beta-OG (0.35%) |
|  | Tip 2 RNAP (8 mg/ml), beta-OG (0.35%) |
| RNAP only | Tip 1 Buffer, beta-OG (0.35%) |
|  | Tip 2 RNAP (8 mg/ml), beta-OG (0.35%) |
| Dynamin tubes + GTP mixed | Tip 1 Dynamin tubes |
|  | Tip 2 GTP (2 mM & 4 mM) |
| Dynamin only | Tip 1 Dynamin tubes |
|  | Tip 2 Buffer |

**Table S6:**
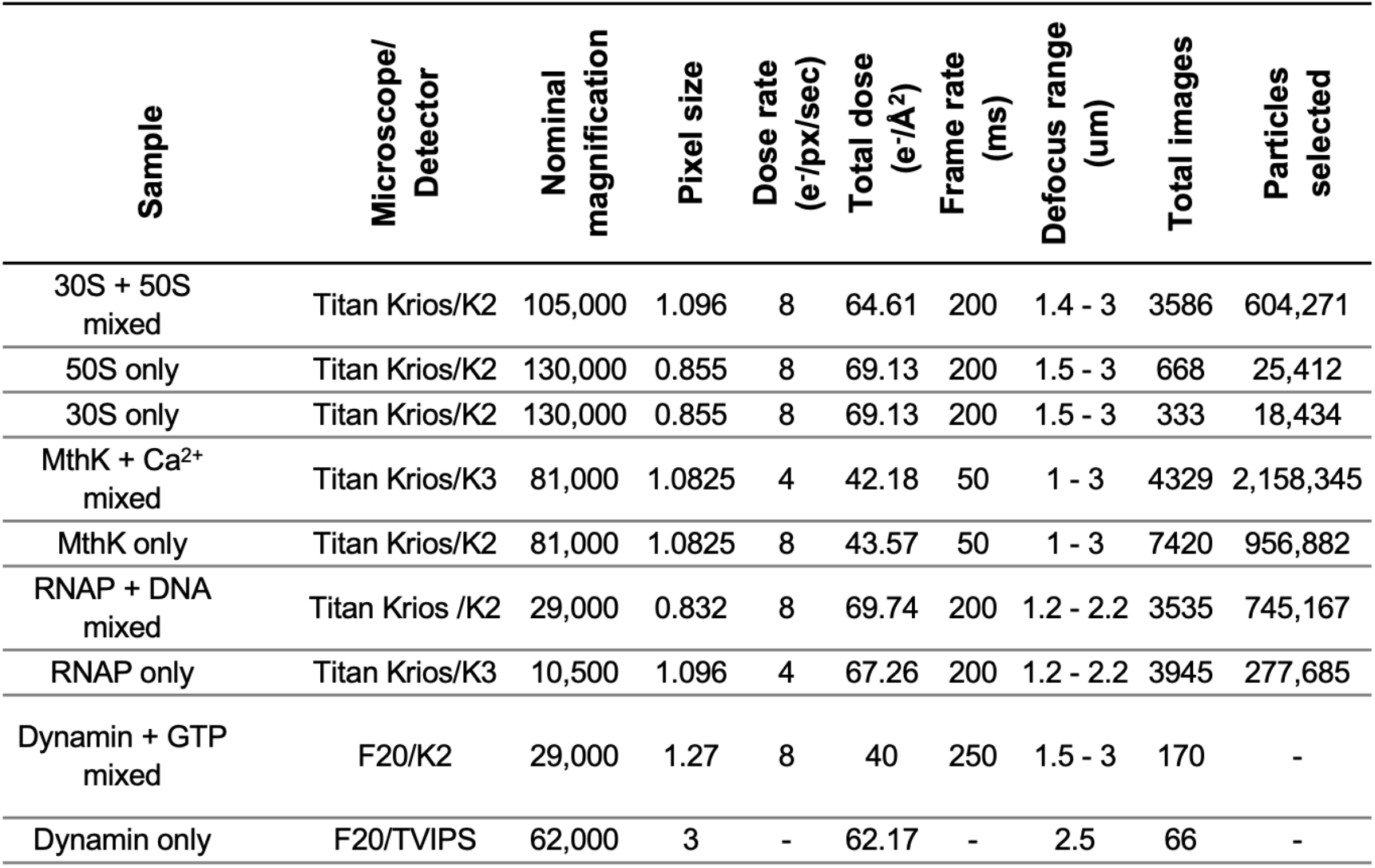
Cryo-EM imaging parameters.

| Sample | Microscope/<br>Detector | Nominal<br>magnification | Pixel size | Dose rate<br>(e-/px/sec) | Total dose<br>(e-/Å<sup>2</sup>) | Frame rate<br>(ms) | Defocus range<br>(μm) | Total images | Particles<br>selected |
| --- | --- | --- | --- | --- | --- | --- | --- | --- | --- |
| 30S + 50S<br>mixed | Titan Krios/K2 | 105,000 | 1.096 | 8 | 64.61 | 200 | 1.4 - 3 | 3586 | 604,271 |
| 50S only | Titan Krios/K2 | 130,000 | 0.855 | 8 | 69.13 | 200 | 1.5 - 3 | 668 | 25,412 |
| 30S only | Titan Krios/K2 | 130,000 | 0.855 | 8 | 69.13 | 200 | 1.5 - 3 | 333 | 18,434 |
| MthK + Ca<sup>2+</sup><br>mixed | Titan Krios/K3 | 81,000 | 1.0825 | 4 | 42.18 | 50 | 1 - 3 | 4329 | 2,158,345 |
| MthK only | Titan Krios/K2 | 81,000 | 1.0825 | 8 | 43.57 | 50 | 1 - 3 | 7420 | 956,882 |
| RNAP + DNA<br>mixed | Titan Krios /K2 | 29,000 | 0.832 | 8 | 69.74 | 200 | 1.2 - 2.2 | 3535 | 745,167 |
| RNAP only | Titan Krios/K3 | 10,500 | 1.096 | 4 | 67.26 | 200 | 1.2 - 2.2 | 3945 | 277,685 |
| Dynamin + GTP<br>mixed | F20/K2 | 29,000 | 1.27 | 8 | 40 | 250 | 1.5 - 3 | 170 | - |
| Dynamin only | F20/TVIPS | 62,000 | 3 | - | 62.17 | - | 2.5 | 66 | - |

## Acknowledgements

We are grateful to the staff of the Simons Electron Microscopy Center at the New York Structural Biology Center for help and technical support. We thank Israel Fernandez and Bridget Huang for kindly providing the ribosome subunits. We thank H. He (NIDDK, NIH) for cryo-EM data collection, NIDDK EM Core Facility, and the Hinshaw lab for critical comments. The work presented here was conducted at the National Resource for Automated Molecular Microscopy located at the New York Structural Biology Center, supported by grants from the NIH (GM103310, OD019994) and the Simons Foundation (SF349247) and the NIDDK NIH Intramural Research Program. Other grant support includes NIH/NIGMS R35 GM118130 (to S.A.D) and NIH/RO1GM088352 (to C.M.N.).

## Data Availability Statement

The data that support the findings of this study are available from the corresponding author upon request.

## References

1 Frank, J. Time-resolved cryo-electron microscopy: Recent progress. J Struct Biol 200, 303–306, doi:10.1016/j.jsb.2017.06.005 (2017).

2 Mulder, A. M. et al. Visualizing ribosome biogenesis: parallel assembly pathways for the 30S subunit. Science 330, 673–677, doi:10.1126/science.1193220 (2010).

3 Sashital, D. G. et al. A combined quantitative mass spectrometry and electron microscopy analysis of ribosomal 30S subunit assembly in E. coli. eLife 3, doi:10.7554/eLife.04491 (2014).

4 Berriman, J. & Unwin, N. Analysis of transient structures by cryo-microscopy combined with rapid mixing of spray droplets. Ultramicroscopy 56, 241–252, doi:10.1016/0304-3991(94)90012-4 (1994).

5 Unwin, N. Acetylcholine receptor channel imaged in the open state. Nature 373, 37–43, doi:10.1038/373037a0 (1995).

6 White, H. D., Walker, M. L. & Trinick, J. A computer-controlled spraying-freezing apparatus for millisecond time-resolution electron cryomicroscopy. J Struct Biol 121, 306–313, doi:10.1006/jsbi.1998.3968 (1998).

7 Unwin, N. & Fujiyoshi, Y. Gating movement of acetylcholine receptor caught by plunge-freezing. J Mol Biol 422, 617–634, doi:10.1016/j.jmb.2012.07.010 (2012).

8 Subramaniam, S. & Henderson, R. Electron crystallography of bacteriorhodopsin with millisecond time resolution. J Struct Biol 128, 19–25, doi:10.1006/jsbi.1999.4178 (1999).

9 Lu, Z. et al. Gas-Assisted Annular Microsprayer for Sample Preparation for Time-Resolved Cryo-Electron Microscopy. J Micromech Microeng 24, 115001, doi:10.1088/0960-1317/24/11/115001 (2014).

10 Lu, Z. et al. Monolithic microfluidic mixing-spraying devices for time-resolved cryo-electron microscopy. J Struct Biol 168, 388–395, doi:10.1016/j.jsb.2009.08.004 (2009).

11 White, H. D., Thirumurugan, K., Walker, M. L. & Trinick, J. A second generation apparatus for time-resolved electron cryo-microscopy using stepper motors and electrospray. J Struct Biol 144, 246–252, doi:10.1016/j.jsb.2003.09.027 (2003).

12 Lu, Z. et al. Passive Microfluidic device for Sub Millisecond Mixing. Sens Actuators B Chem 144, 301–309, doi:10.1016/j.snb.2009.10.036 (2010).

13 Shaikh, T. R. et al. Initial bridges between two ribosomal subunits are formed within 9.4 milliseconds, as studied by time-resolved cryo-EM. Proc Natl Acad Sci U S A 111, 9822–9827, doi:10.1073/pnas.1406744111 (2014).

14 Chen, B. et al. Structural dynamics of ribosome subunit association studied by mixing-spraying time-resolved cryogenic electron microscopy. Structure 23, 1097–1105, doi:10.1016/j.str.2015.04.007 (2015).

15 Fu, Z. et al. Key Intermediates in Ribosome Recycling Visualized by Time-Resolved Cryoelectron Microscopy. Structure 24, 2092–2101, doi:10.1016/j.str.2016.09.014 (2016).

16 Kaledhonkar, S. et al. Late steps in bacterial translation initiation visualized using time-resolved cryo-EM. Nature 570, 400–404, doi:10.1038/s41586-019-1249-5 (2019).

17 Kontziampasis, D. et al. A cryo-EM grid preparation device for time-resolved structural studies. IUCrJ 6, 1024–1031, doi:10.1107/S2052252519011345 (2019).

18 Jain, T., Sheehan, P., Crum, J., Carragher, B. & Potter, C. S. Spotiton: a prototype for an integrated inkjet dispense and vitrification system for cryo-TEM. J Struct Biol 179, 68–75, doi:10.1016/j.jsb.2012.04.020 (2012).

19 Dandey, V. P. et al. Spotiton: New features and applications. J Struct Biol 202, 161–169, doi:10.1016/j.jsb.2018.01.002 (2018).

20 Wei, H. et al. Optimizing “self-wicking” nanowire grids. J Struct Biol 202, 170–174, doi:10.1016/j.jsb.2018.01.001 (2018).

21 Scapin, G. et al. Structure of the insulin receptor-insulin complex by single-particle cryo-EM analysis. Nature 556, 122–125, doi:10.1038/nature26153 (2018).

22 Xu, K. et al. Epitope-based vaccine design yields fusion peptide-directed antibodies that neutralize diverse strains of HIV-1. Nat Med 24, 857–867, doi:10.1038/s41591-018-0042-6 (2018).

23 Zhang, Z. et al. Ensemble cryoEM elucidates the mechanism of insulin capture and degradation by human insulin degrading enzyme. eLife 7, doi:10.7554/eLife.33572 (2018).

24 Han, H. et al. Structure of Vps4 with circular peptides and implications for translocation of two polypeptide chains by AAA+ ATPases. eLife 8, e44071, doi:10.7554/eLife.44071 (2019).

25 Koh, F. et al. The structure of a 15-stranded actin-like filament from Clostridium botulinum. Nature Communications 10, 2856, doi:10.1038/s41467-019-10779-9 (2019).

26 Liu, Y. et al. FACT caught in the act of manipulating the nucleosome. Nature 577, 426–431, doi:10.1038/s41586-019-1820-0 (2020).

27 Wu, J. L. Y., Tellkamp, F., Khajehpour, M., Robertson, W. D. & Miller, R. J. D. Rapid mixing of colliding picoliter liquid droplets delivered through-space from piezoelectric-actuated pipettes characterized by time-resolved fluorescence monitoring. Rev Sci Instrum 90, 055109, doi:10.1063/1.5050270 (2019).

28 Jiang, Y. et al. The open pore conformation of potassium channels. Nature 417, 523–526 (2002).

29 Jiang, Y. et al. Crystal structure and mechanism of a calcium-gated potassium channel. Nature 417, 515–522 (2002).

30 Zadek, B. & Nimigean, C. M. Calcium-dependent gating of MthK, a prokaryotic potassium channel. J Gen Physiol 127, 673–685 (2006).

31 Ye, S., Li, Y., Chen, L. & Jiang, Y. Crystal structures of a ligand-free MthK gating ring: insights into the ligand gating mechanism of K+ channels. Cell 126, 1161–1173 (2006).

32 Ruff, E. F., Record, M. T., Jr. & Artsimovitch, I. Initial events in bacterial transcription initiation. Biomolecules 5, 1035–1062, doi:10.3390/biom5021035 (2015).

33 Mazumder, A. & Kapanidis, A. N. Recent Advances in Understanding sigma70-Dependent Transcription Initiation Mechanisms. J Mol Biol 431, 3947–3959, doi:10.1016/j.jmb.2019.04.046 (2019).

34 Saecker, R. M., Record, M. T., Jr. & Dehaseth, P. L. Mechanism of bacterial transcription initiation: RNA polymerase - promoter binding, isomerization to initiation-competent open complexes, and initiation of RNA synthesis. J Mol Biol 412, 754–771, doi:10.1016/j.jmb.2011.01.018 (2011).

35 Sundborger, A. C. et al. A dynamin mutant defines a superconstricted prefission state. Cell Rep 8, 734–742, doi:10.1016/j.celrep.2014.06.054 (2014).

36 Kong, L. et al. Cryo-EM of the dynamin polymer assembled on lipid membrane. Nature 560, 258–262, doi:10.1038/s41586-018-0378-6 (2018).

37 Johansson, M., Bouakaz, E., Lovmar, M. & Ehrenberg, M. The kinetics of ribosomal peptidyl transfer revisited. Mol Cell 30, 589–598, doi:10.1016/j.molcel.2008.04.010 (2008).

38 Chen, J. et al. E. coli TraR allosterically regulates transcription initiation by altering RNA polymerase conformation. eLife 8, doi:10.7554/eLife.49375 (2019).

39 Posson, D. J., Rusinova, R., Andersen, O. S. & Nimigean, C. M. Calcium ions open a selectivity filter gate during activation of the MthK potassium channel. Nat Commun 6, 8342, doi:10.1038/ncomms9342 (2015).

40 Suloway, C. et al. Automated molecular microscopy: the new Leginon system. J Struct Biol 151, 41–60, doi:10.1016/j.jsb.2005.03.010 (2005).

41 Zheng, S. Q. et al. MotionCor2: anisotropic correction of beam-induced motion for improved cryo-electron microscopy. Nat Methods 14, 331–332, doi:10.1038/nmeth.4193 (2017).

42 Rohou, A. & Grigorieff, N. CTFFIND4: Fast and accurate defocus estimation from electron micrographs. J Struct Biol 192, 216–221, doi:10.1016/j.jsb.2015.08.008 (2015).

43 Sorzano, C. O. et al. A clustering approach to multireference alignment of single-particle projections in electron microscopy. J Struct Biol 171, 197–206, doi:10.1016/j.jsb.2010.03.011 (2010).

44 Lander, G. C. et al. Appion: an integrated, database-driven pipeline to facilitate EM image processing. J Struct Biol 166, 95–102 (2009).

45 Roseman, A. M. FindEM--a fast, efficient program for automatic selection of particles from electron micrographs. J Struct Biol 145, 91–99, doi:10.1016/j.jsb.2003.11.007 (2004).

46 Schindelin, J. et al. Fiji: an open-source platform for biological-image analysis. Nat Methods 9, 676–682, doi:10.1038/nmeth.2019 (2012).

47 Fan C., Sukomon N., Flood E., Rheinberger J., Allen T.W., and Nimigean C.M. Ball-and-chain inactivation in a calcium-gated potassium channel. Nature. https://doi.org/10.1038/s41586-020-2116-0 (2020)

48 Scheres SH. A Bayesian view on cryo-EM structure determination. J Mol Biol. 2012;415(2):406–18. Epub 2011/11/22. doi: 10.1016/j.jmb.2011.11.010. PubMed PMID: 22100448; PMCID: PMC3314964.

49 Punjani A, Rubinstein JL, Fleet DJ, Brubaker MA. cryoSPARC: algorithms for rapid unsupervised cryo-EM structure determination. Nat Methods. 2017;14(3):290–6. Epub 2017/02/07. doi: 10.1038/nmeth.4169. PubMed PMID: 28165473.

